# Disease-causing MFN2 mutants impair mitochondrial fission dynamics by distinct DRP1 dysregulation

**DOI:** 10.1101/2025.04.03.647131

**Authors:** Daniel Lagos, Pamela R. de Santiago, Nicolás Pérez, Benjamín Cartes-Saavedra, Josefa Vial-Brizzi, Oliver Podmanicky, Rita Horvath, Verónica Eisner

## Abstract

Mitochondria undergo fusion and fission. While DRP1 regulates fission, fusion is controlled by OPA1, MFN1, and MFN2. The balance between these processes and the crosstalk between machineries remains poorly understood. MFN2 mutations cause Charcot-Marie-Tooth disease type 2A (CMT2A), affecting mitochondrial fusion and morphology. However, their role in fission is unclear.

Using skin fibroblasts from CMT2A patients (L248H and M376V MFN2 mutations) and wild-type mouse embryonic fibroblasts expressing these variants we studied how MFN2 mutations impact mitochondrial dynamics beyond fusion. We analysed mitochondrial morphology and dynamics, by live-cell confocal microscopy, and tested fusion/fission protein levels, oxygen consumption rate (OCR), extracellular acidification rate (ECAR), and oxidative phosphorylation complex subunits.

MFN2 mutations impaired mitochondrial fusion and displayed distinct effects on fission and cellular metabolism. L248H-expressing cells showed hyper-elongated mitochondria, impaired fission, and increased OCR, while, M376V cells exhibited fragmentation, enhanced fission, and elevated ECAR. These effects correlated with differential Drp1 phosphorylation.

Our findings demonstrate that MFN2 mutants differentially influences fission and metabolism, highlighting the need to consider these effects in therapies aimed at modulating mitochondrial dynamics.

## Introduction

Mitochondria serve as the energetic hub within the cell. To properly accomplish their functions mitochondria can respond to extra and intracellular stimuli by modulating their complex morphology, undergoing fusion and fission processes ^1^. Mitochondrial fusion involves sequential steps. It starts with the interaction of mitochondria through the homo or heterodimers of the outer mitochondrial membrane (OMM) GTPases Mitofusin-1 and 2 (MFN1/2) ^2,3^, allowing adjacent membrane close positioning and fusing as a result of conformational changes dependent on GTP hydrolysis ^4^. Following OMM fusion, the inner mitochondrial membrane (IMM) GTPase OPA1 mediates the fusion of its compartment, leading to the subsequent mixing of mitochondrial matrix contents ^5^. Fusion results in a larger mitochondrion with mixed mitochondrial components ^6^. Mitochondrial fission, instead, allows the division of one mitochondrion into two individual units. In mammals, the GTPase Dynamin-related protein 1 (DRP1) is constantly recruited to the mitochondria from the cytoplasm by OMM resident proteins such as Mitochondrial fission factor (MFF) ^7^ and Mitochondrial elongation factor (MIEF1/2) ^8^. Once on the OMM, the maturation of DRP1 oligomers in fission-productive oligomers relies on their phosphorylation status ^9,10^ and in polymerization of actin filaments in the endoplasmic reticulum (ER)-mitochondria contact sites ^11^. Mitochondrial fission allows the segregation of damaged mitochondria through mitophagy ^12^, exchange of mitochondrial DNA (mtDNA) to the nascent mitochondria as replication occurs in fission sites ^13^, inheritance of mitochondria to daughter cells during cell division ^14^, and localization of mitochondria at neuromuscular junctions ^15^. Dysregulation of fusion or fission by loss of the protein that controls these processes results in pathological consequences that range from mtDNA instability, abnormal mitochondrial morphology, and metabolic collapse to severe diseases or premature death. Thus, a precise fusion and fission balance is vital for organelle and cellular physiology.

Mitochondrial fusion and fission are opposite but interconnected phenomena and require precise cross-regulation. It has been suggested that both occur in the ER-mitochondria contact sites, where the fusion and fission proteins converge ^16,17^. This overlap determines a complex interplay between fusion and fission proteins, where protein-protein interaction seems to play a fundamental role in maintaining the balance. Overexpression of a cytoplasmic form of Mfn2 elongates mitochondria, interfering with the fission machinery, specifically by Drp1-Mfn2 physical interaction and its cytoplasm sequestration ^18^. Whether this interaction occurs in the OMM is not known. Mitochondrial fission 1 protein (FIS1) modulates mitochondrial fission through functional interaction with proteins MFN1/2 and OPA1, inhibiting their GTPase activity and, therefore, mitochondrial fusion ^19^. In addition, MIEF1 interferes with the fusion process by interacting with MFN1/2 and competitively decreases their interaction with FIS1, enhancing mitochondrial fusion ^20^. Despite these mechanistic pieces of evidence, there is a lack of studies addressing fusion and fission dynamics in pathological models that affect mitochondrial fusion or fission proteins.

Mutations in *MFN2* lead to the development of CMT2A, the most common form of axonal CMT, characterized by distal motor and sensory impairment ^21^. Heterozygous-missense mutations are the most frequent type of genetic defect driving CMT2A ^22^. The mechanism underlying the axonal degeneration is not entirely understood. It seems to rely on a complex interplay between the processes in which MFN2 participates, such as mitochondrial transport ^23–25^, ER-mitochondria interaction ^26,27^, and fusion ^28,29^. In addition, nearly 136 pathogenic *MFN2* single nucleotide variants have been reported (ClinVar Database) ^30^, resulting in a broad phenotype spectrum at cellular and clinical levels ^31^. Hence, it is necessary to continue exploring *MFN2* mutations and their effects on mitochondrial bioenergetics.

MFN2-R364W overexpression causes mitochondrial hyperelongation involving Drp1 dysregulation ^32,33^. In patients, this mutation produces an early-onset presentation of CMT2A and severe peripheral neuropathy with optic atrophy ^31,34^. Hyperelongation of mitochondria prevents excessive mitophagy and fission-mediated apoptosis ^35^. However, sustained mitochondrial hyperelongation impairs its trafficking, depleting the mitochondria from the neuromuscular-junction ^32^, and compromises its quality control ^36^. A mutation in DRP1 that prevents its oligomerization has been associated with a severe, fatal disease in a child, supporting the importance of fission and the deleterious effects of chronic hyperelongation ^37,38^. Nevertheless, whether mitochondrial fission disbalance contributes to the pathological mechanism of CTM2A is unknown.

In this work, we comprehensively analyzed the effects of different *MFN2* mutations on mitochondrial fusion and fission dynamics. We used patient-derived fibroblasts and MEF cell lines acutely or chronically expressing the severe *MFN2* c.742T>A (MFN2-L248H) ^31,39^ or the milder variant c.1126A>G (MFN2-M376V) ^40,41^. We found that both mutants decreased the fusion frequency but displayed different mitochondrial phenotypes. While M376V cells showed a fragmented mitochondrial phenotype, L248H mitochondria were hyperelongated, with a decrease in fission frequency and longer fission time, associated with an increase in pDRP1-S637 phosphorylation, severe mitochondrial cristae disruption, and increased oxygen consumption. We propose that some MFN2 mutations may alter not only fusion but also fission dynamics, leading to hyperelongation, potentially underlying severe forms of CMT2A.

## Materials and methods

### Cell culture

Experiments were performed in skin fibroblasts derived from CMT2A patients and control individuals or MEF cells. Patient-derived cells were provided by the Newcastle Research Biobank for Rare and Neuromuscular Diseases, based on a Material Transference Agreement with Pontificia Universidad Católica de Chile. All the cells were cultured in high glucose Dulbecco-Eagle modified medium containing sodium pyruvate (DMEM) and supplemented with 10% of fetal bovine serum (FBS), 2mM glutamine, and 100 U/ml penicillin, 100 μg/mL streptomycin in humidified air (5% CO2) at 37°C. The patient’s fibroblasts were used between passages 3-10. All cells were tested for mycoplasma by qPCR using the described primers ^42^. The cells were cultured following the institutional biosafety and medical ethics protocols.

### Western blot analysis

We cultured the cells to 70-80% of confluence, harvested and frozen at −80 °C. After thawing, a whole cell lysate was generated using RIPA buffer supplemented with protease and phosphatase inhibitors; 30 μg of total protein extracts were loaded into 10% SDS-PAGE gel and transferred to PVDF membranes. Membranes were blocked with 5% milk in 0.1% TBS-Tween for 1 hour at room temperature, followed by an overnight incubation with primary antibody diluted in 5% milk or 3% BSA in 0.1% TBS-Tween (Supplementary Table 1. Antibodies). The following HRP-conjugated secondary antibodies were prepared in 5% milk and 0.1% TBS-Tween. The chemiluminescent reaction was visualized using HRP-enhanced substrates (ECL, SuperSignal West Dura or SuperSignal West Femto, Thermo Fisher). Densitometry was performed using the open-source software Fiji ^43^.

### Cell transfection

Cells were plated on glass coverslips and transfected with 1 μg of each specific construct. Plasmids (Supplementary Table 2. Plasmids) were delivered using Opti-MEM and Lipofectamine 2000 or 3000 (Thermo Fisher) according to the commercial protocol. The cells were cultured 24-48 hours after transfection for optimal expression.

### Mitochondrial dynamics studies by confocal microscopy

Image acquisition of human fibroblasts and MEF cells was performed in a 0.25% BSA extracellular medium (ECM; 121 mM NaCl, 5 mM NaHCO3, 4.7 mM KCl, 1.2 mM KH2PO4, 1.2 mM MgSO4, 2 mM CaCl2, 10 mM glucose, and 10 mM Na-Hepes, pH 7.4), at 37 °C. Mitochondrial matrix fusion dynamics of CMT2A-derived fibroblast and MEFs was studied at a laser-scanning microscope Nikon Eclipse C2 or Nikon TimeLapse, using a 63X/1.4 ApoPlan objective, located at the Advanced Microscopy Facility of the Pontificia Universidad Católica de Chile (UMA-UC). Time series experiments of mtPA-GFP and mtDsRed were executed by excitation with 488 nm and 560 nm laser lines, respectively, every 3 s, for a total of 7 min. A region of interest (ROI) of 5 μm x 5 μm was selected to photoactivate mtPA-GFP with 405 nm the laser line for 10 s. Mitochondrial fusion events were quantified by evaluating the areas around the photoconverted ROIs as previously described ^44^. Briefly, we identify the close apposition of a photoactivated mitochondrion (donor) and a non-photoactivated one (acceptor), followed by the quantic increase in the GFP fluorescence in the acceptor mitochondrion and, concomitantly, a quantic decrease in mtPA-GFP in the donor mitochondrion. The fusion events frequency was analyzed as previously described, considering events/min/3ROI.

### Mitochondrial fission frequency

Mitochondrial fission frequency was quantified as previously reported ^11^. Briefly, the patient’s cells transfected with mtDsRed were imaged in Nikon Eclipse C2, using a 63X/1.4 ApoPlan objective every 3 s for 7 min. Stable MEF cells expressing mitochondria-targeted Blue Fluorescent Protein (mtBFP) and Yellow Fluorescent Protein-Drp1 (EYFP-Drp1) were imaged by high-resolution confocal microscopy Zeiss 880 with an Airyscan detector at UMA-UC, using the 408 nm, and the 488 nm laser lines, respectively. Time series were generated in the “fast mode” by utilizing 16 of the 32 available detectors. We registered one z-stack of 5 slices of 0.19 µm steps every 4 s (0.25 Hz) for 7 minutes. The Zeiss Zen Blue software further processed raw data. Airyscan data were analyzed as a Z-projection of 5 slices.

Four to five ROIs were selected in each cell at sub-regions where a discrete mitochondrion was resolvable and remained in the focal plane. Mitochondrial length varies depending on cell type. In human-skin fibroblast, it has been reported near to 6.5 μm ^45^. As mitochondrial length influences the rate of fission, thus short mitochondria have been shown to have a low fission rate; for this analysis, we considered mitochondria longer than 2 µm ^46^. ROIs were manually scanned frame-by-frame, searching for mitochondrial fission events. The separation of two daughter mitochondria was confirmed when the fluorescence intensity between them was comparable with the background. Finally, mitochondrial length was manually determined. Data were plotted as the number of fission events per minute per length of mitochondria. The average fission frequency from 4-5 ROIs per cell was graphed. We also measured the number of constrictions per length and the lag time, which we defined as the period between the first constriction observation and the actual mitochondrial division. For EYFP-Drp1 images, we applied a threshold to confirm the presence of Drp1 at fission sites (Fig. S4A).

### Transmission electron microscopy (TEM)

To evaluate mitochondrial ultrastructure, 8×10^5^ cell pellets were fixed with 2.5% glutaraldehyde. Staining was performed as previously described ^47^. Ultrathin sections were imaged in the transmission electron microscope Philips Tecnai 12 at 80 kV at UMA-UC. Area and perimeter were quantified using the software Fiji. Semiquantitative characterization of mitochondria cristae was made using the following classifiers: Structured, mitochondria with regular and deeply folded cristae; Empty, mitochondria without cristae; Irregular, mitochondria with short and discontinuous folding, showing matrix with clear or empty areas; Aberrant, mitochondria displaying empty regions along with structured or irregular zones.

### Stable cell line generation

#### MFN2-Flag cloning

The human MFN2 gene was PCR amplified from pECFP-C1-MFN2 using cloning primers (Supplementary Table 3. Primers and probes), adding BamHI cutting site at 3’ and Flag sequence followed by EcoRI cutting site at 5’ of the MFN2 coding sequence. The PCR product was double digested with EcoRI and BamHI High-fidelity enzymes in rCutsmartTM Buffer (NEB), generating cohesive ends. The product was cloned into a third-generation lentiviral plasmid hUbC-driven EGFP, FUGW. E

The GFP sequence was replaced by the MFN2-Flag sequence. Ligation was performed with T4 DNA ligase overnight at 4°C. Chemically competent *Escherichia coli XL10* was transformed with ligation products using standard heat shock transformation protocol. Bacteria were grown overnight, and plasmids were purified (Macherey-Nagel Midiprep kit). MEF cells were transfected with the purified MFN2-Flag plasmid, and flag expression was tested.

#### Mutant MFN2-Flag generation

Mutagenic MFN2-Flag variants were designed using the Q5 Site-Directed Mutagenesis Kit (NEB) following the manufacturer’s protocol. Products were sequenced using the ABI PRISM 3500 xl Applied Biosystems at Pontificia Universidad Católica de Chile. Monoclonal stable cell lines generation: MFN2-Flag and mutants were transfected in WT MEFs. Three days after transfection, cells were selected with 400 ug/ml Bleomycin antibiotic (Zeocin, Thermo Fisher). Cells were maintained in Zeocin-supplemented media. Six days after selection individual colonies were observed, manually picked, and transferred to 96 well plates for further culture. Mitochondrial morphology and MFN2-Flag expression were tested to select working clones (Fig. S2A).

### Quantification of mtDNA levels

We performed the relative quantification of mtDNA levels in stable MEFs expressing empty vector, WT, or mutant MFN2. Total DNA was extracted using the PureLink genomic DNA Minikit. Amplification of the *Mus musculus* mitochondrial ND5 (mt-ND5) gene and the nuclear-encoded ACTB (β-actin) was performed using TaqMan-qPCR (Supplementary Table 3. Primers and probes). Relative mtDNA amount was calculated as the Ct value difference of β-actin and mt-ND5, where Delta Ct (ΔCt) equals the sample Ct of the mt-ND5 subtracted from the sample Ct of the nuclear reference gene.

### Long-range PCR

Long-range PCR assays were performed to study mtDNA deletions in stable MEFs. Two fragments covering the major and minor arc of the *Mus musculus* mtDNA were amplified using specific mtDNA primers (Supplementary Table 3. Primers and probes). PrimeSTAR GXL polymerase (Takara Bio) and 100 ng of template DNA were used. An overnight reaction of 30 cycles was performed. Reaction products were run on 0.7% agarose-TAE gel for 1.5 h and visualized by UV transillumination.

### Mitochondrial oxygen consumption and extracellular acidification assay

Oxygen consumption rate (OCR) and extracellular acidification rate (ECAR) of stable MEFs were determined using the XF96 Extracellular Flux Analyzer (Seahorse Bioscience, Agilent Technologies). Cells were seeded in a 96-well microplate (Seahorse Bioscience) 24 h before the experiment using 80 μl of medium. Sequential addition of oligomycin 1 μM, carbonyl cyanide-p-trifluoromethoxy-phenylhydrazone (FCCP) 1 μM, rotenone 1μM, and antimycin + rotenone 1 μM allowed the determination of oxygen consumption rates attributable to basal respiration, proton leakage, maximal respiratory capacity, and non-mitochondrial respiration. ATP-linked respiration and reserve capacity were also calculated. Basal ECAR and Glycolytic capacity, calculated as the difference between ECAR following the injection of 1 μm oligomycin, and the basal ECAR reading was reported. OCR and ECAR levels were normalized to cell number per well estimated by counting fluorescently labelled nuclei from images captured by Cytation 5. Hoechst was included in the last injection in the XF analysis protocol, with the rotenone/antimycin A injection.

### Mitochondrial membrane potential

Cells were plated on glass coverslips, washed with ECM, and loaded for 10 min at room temperature with 10 nM Tetramethyl rhodamine methyl ester (TMRE) in non-quenching mode. Cells were imaged at the Nikon Eclipse Ti microscope with the following configuration: ex. 540 nm—em. 620/60 nm, one image every 1.5s. FCCP 5 µM was used to induce loss of mitochondrial membrane potential. Resting membrane potential was calculated as ΔF_basal_-F_FCCP_ using absolute fluorescence values.

### Statistical analysis

The statistical analysis was carried out using GraphPad Prism 8 Software. For multiple comparisons, One-way ANOVA followed by Dunnett’s or Sidak’s multiple comparison tests was used to determine the significance of normally distributed data or Two-way ANOVA followed by multiple t-test comparison was used to assess significance. For non-parametric multiple comparisons, a Kruskal-Wallis test followed by Dunn’s multiple comparison tests was used to determine significance. In all cases, data not indicated as significant should be considered not statistically different.

## Results

### Cells carrying MFN2-mutations L248H and M376V display different mitochondrial shape and cristae alterations

We studied skin-derived fibroblasts from CMT2A-patients carrying the heterozygous mutations c.742T>A (*MFN2*-L248H) and c.1126A>G (*MFN2-*M376V), affecting the GTPase and HR1 domains, respectively (Fig. 1A). At the moment of biopsy, the patients manifested differential phenotypes, with a more severe and early onset presentation in L248H and milder and later onset disease in the patient with M376V mutation (Table I). To evaluate whether these phenotypic differences correlate with differences at the cellular level, we studied the mitochondrial ultrastructure using TEM. Cristae ultrastructure was classified as structured, empty, irregular, or aberrant cristae. Representative images of each classifier are presented in Fig. 1B. We found that cells carrying the L248H mutation exhibited a pronounced decrease in the abundance of mitochondria with structured cristae (5%) and an increase in the percentage of mitochondria devoid of cristae or empty mitochondria (45%). Mitochondria from M376V-fibroblasts showed a lesser extent of cristae alteration, displaying a non-significant decrease in structured (32%) and increase in aberrant (32%) mitochondria, compared to control cells (structured 46%, empty 4%, irregular 35% and aberrant 15%) (Fig. 1B-C). These data suggest distinct MFN2 mutant-dependent cristae remodelling defects.

**Figure 1.**
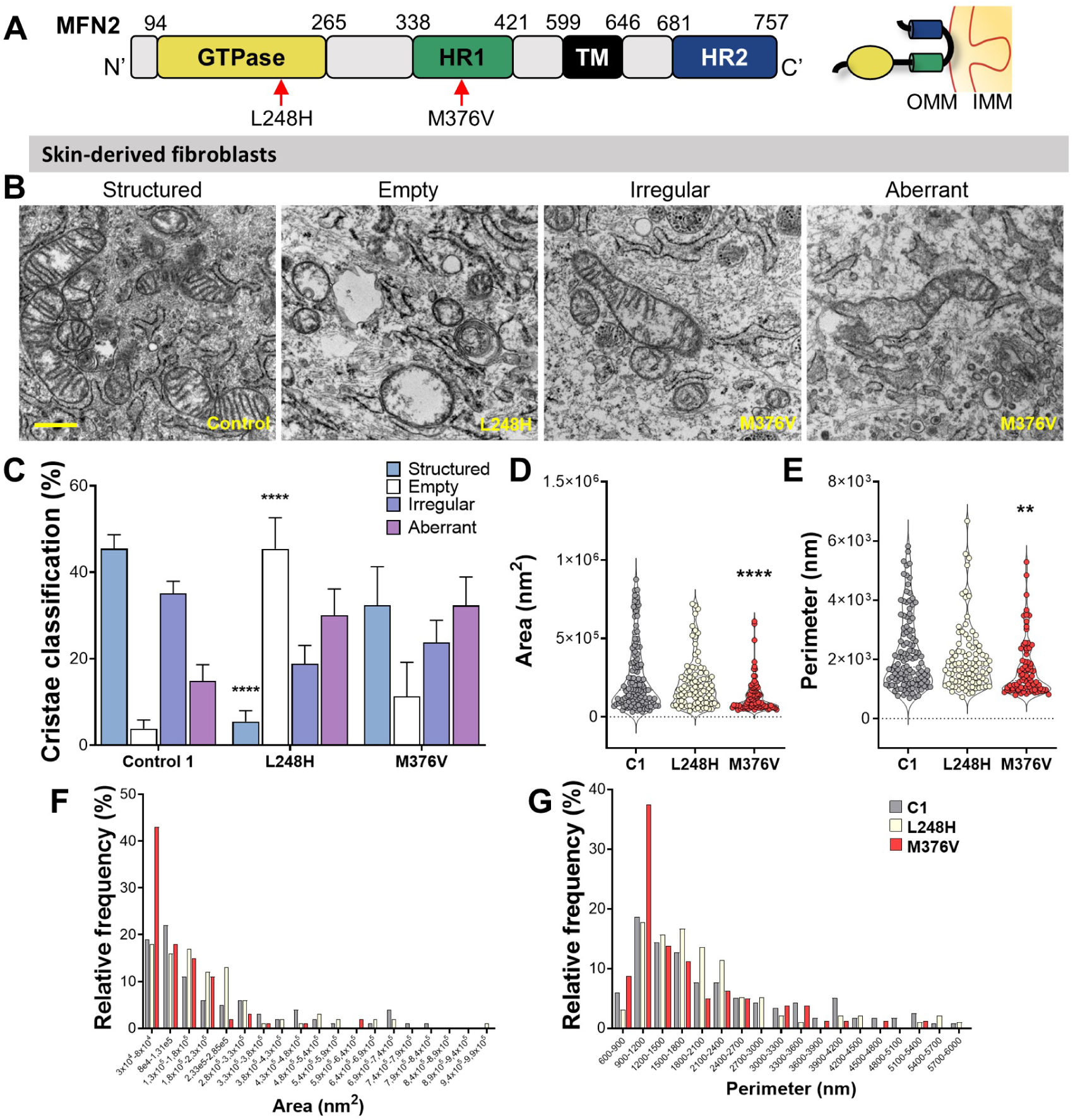
Mitochondrial ultrastructure of CMT2A-derived fibroblasts is differentially affected. (**A**) Functional domains of MFN2: GTPase domain (yellow); Heptad Repeat 1 (HR1, green); transmembrane domain (TM, black); and Heptad Repeat 2 (HR2, blue). MFN2 is located in the OMM, with the GTPase, HR1, and HR2 domains exposed to the cytoplasm. (**B**) Representative TEM images of mitochondrial cristae classification from CMT2A-derived fibroblasts. Scale bar = 500 nm. (**C**) Quantification of cristae descriptors. (**D**) Mitochondrial area and (**E**) perimeter quantification. (**F**) mitochondrial area and (**G**) perimeter frequency distribution. Control 1 n = 2/11/118 (preparations/cells/mitochondria), L248H n = 3/18/96, M376V n = 2/13/80. Data are mean ± SEM of 3 independent experiments. \*\**P*<0.01, \*\*\*\**P*<0.0001 *vs* Control 1.

Quantifying mitochondrial area and perimeter showed that fibroblasts carrying the L248H mutant (Fig. 1D-E) presented comparable area and perimeter. However, we observed a relative increase in the frequency of mitochondria with large regions (1,8×10^5^– 2,8×10^5^ nm [Fig. F]) and perimeter (1200 −2400 nm [Fig. 1G]). The cells carrying M376V mutant presented smaller mitochondria with decreased total area and perimeter (Fig. 1D-E), with nearly 40% of mitochondria grouped in the smallest ranges of area (3×10^4^ – 8×10^4^ nm [Fig. F]) and perimeter (900-1200 nm [Fig. G]). Thus, the ultrastructure analysis of CMT2A patient-derived cells suggests a differential mechanistic dysregulation for each studied MFN2 mutant.

### MFN2-mutations L248H and M376V differentially alter mitochondrial morphology and decrease mitochondrial fusion frequency

#### Studies in CMT2A patient-derived fibroblasts

To determine whether the differences in mitochondrial morphology observed in TEM results arise from a defect in mitochondrial fusion activity related to MFN2’s role in OMM fusion, we co-transfected patient-derived fibroblasts with the mitochondrial-targeted fluorescent proteins mtDsRed and mtPA-GFP (Fig. 2A). Mitochondrial morphology was determined and classified using the mtDsRed signal. We found a high percentage of L248H-carrying cells with elongated mitochondria and the absence of fragmentation. In contrast, the M376V-carrying fibroblasts showed mainly a fragmented phenotype (Fig. 2A-B), confirming our previous observations.

**Figure 2.**
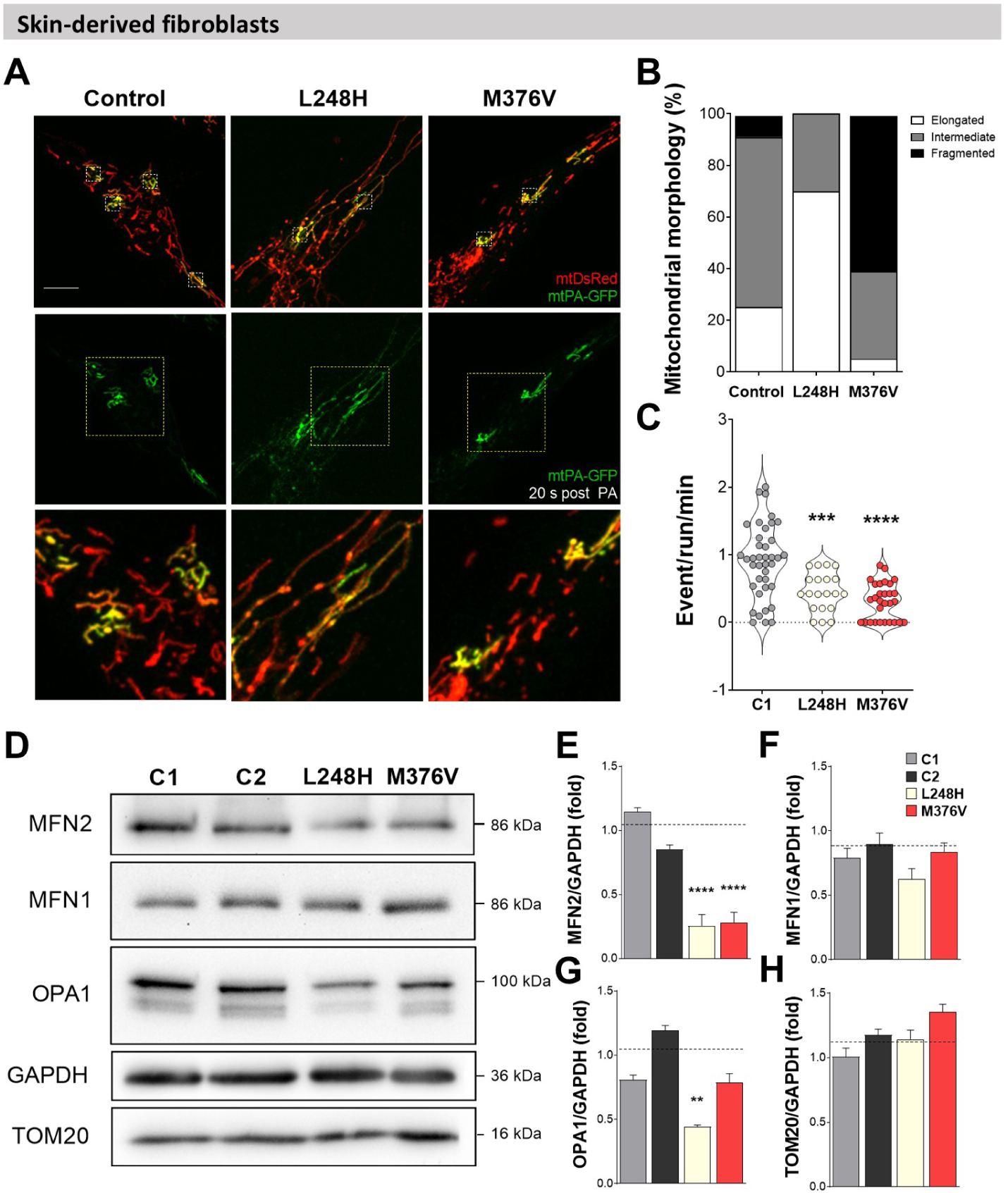
MFN2 mutants L248H and M376V decrease fusion frequency and differentially alter mitochondrial morphology in CMT2A patient-derived fibroblasts. (**A**) Control or CMT2A-derived fibroblasts expressing mtDsRed (red) and mtPA-GFP (green). Panels show representative images 20 s after photoactivation of 5 x 5 μm ROIs (white squares, top) by 408 nm laser illumination and acquired upon 488 and 560 nm laser excitation. Photoconverted areas depicting mtPA-GFP diffusion (middle) and insets from the top panel illustrating the mitochondrial morphology (bottom). Scale bar 10 μm. (**B**) Mitochondrial morphology classification and quantification using mtDsRed images in ≥ 30 cells. (**C**) Fusion events frequency (Control = 39 cells, L248H = 20 cells, M376V = 29 cells). (**D**) Representative Western blot from whole protein extracts of controls (C1 and C2) or CMT2A-derived fibroblasts carrying MFN2 mutants. (**E**), (**F**), (**G**), and (**H**) Western blot densitometric quantification. The dashed line represents the average of controls. Data are mean ± SEM of ≥ 3 independent experiments. **P<0.01, ***P<0,001, ****P<0.0001 *vs* Control 1.

Next, we photoconverted the mtPA-GFP and followed the cells for 7 minutes, as described in the methods section. Quantification of mitochondrial fusion frequency showed that L248H and M376V mutants displayed a reduction in fusion frequency despite the differences observed in morphology (Fig. 2C). Immunoblot analysis revealed that the impaired fusion was accompanied by low levels of MFN2 (Fig. 2D-E) with no changes in MFN1 levels (Fig.2D-F) in both cell lines. We also found that cells carrying the mutant L248H showed low levels of OPA1 (Fig. 2D-G), a protein critical on IMM fusion and cristae structure, consistent with the cristae alterations that this mutant shows.

#### Studies in Mfn2^−/−^ background

Because patient’s cells can carry different genetic backgrounds and have been subjected to distinct environmental conditions, we used an isogenic background to evaluate the effect of the MFN2 mutants using *Mfn2* ^−/−^ MEFs. For this purpose, we used the pECFP-C1-MFN2 plasmid and performed point mutations to introduce the MFN2 mutants c.742T>A/ L248H or c.1126A>G/ M376V. As expected, rescue experiments showed that expression of WT-MFN2 rescued the fragmented phenotype of *Mfn2* ^−/−^ cells, increasing the number of cells with an elongated or intermediate mitochondrial network (Fig. S1A-B). Also, the acute expression of L248H promoted the rescue of the mitochondrial morphology comparable to the WT; meanwhile, M376V-expressing cells only showed a partial mitochondria morphology rescue with a high abundance of cells with an intermediate phenotype (Fig. S1A-B). The observed phenotypes differ from those observed in patients’ cells (L248H/elongation and M376/fragmentation); however, as in human fibroblasts, mitochondrial fusion frequency was lower than in the cells expressing MFN2 mutations when compared with WT-MFN2 expression, despite the differences observed at the level of the morphology. Also, the expression of MFN2 mutants was lower than the WT, similar to the observed in the patient cells (Fig. S1D), suggesting that the control of the mitochondrial morphology by MFN2 is not only restricted to the fusion activity and protein levels.

#### Studies in WT background

As our patient cells carry heterozygous mutations for MFN2, implying the co-existence of a WT and a mutant allele, we hypothesized that the phenotypes observed in these cells could be a result of a “dominant-negative” effect as previously proposed for the pathological mechanism of MFN2 mutations in CMT2A ^52 53^. To test this, we acutely expressed WT and the MFN2 mutants in WT MEFs background. Mitochondrial morphology analysis showed that expression of L248H increased the number of cells showing an elongated mitochondrial population compared to overexpression of WT MFN2 (*Elongated frequency*: WT MFN2=28%; L248H=72%), with no cells displaying fragmentation. In contrast, expression of the M376V mutants led to cells with fragmented mitochondria (*Fragmented frequency:* WT MFN2=5%; M376V=58%) (Fig. 3A-B). Analysis of mitochondrial fusion in these cells showed that expression of the two MFN2 mutations had a detrimental effect on the mitochondrial fusion frequency, suggesting that they could be acting as dominant negative (Fig. 3C). Interestingly, cells with no fusion events in the WT background were observed only in those expressing the MFN2 mutations and not during the expression of the WT MFN2 or the mock condition (*# cells with no fusion events*: MFN2=0; L248H=6; M376V=10) (Fig. 3C), suggesting a blockade of the fusion process in the presence of these mutants. However, the fusion inhibition does not explain mitochondrial elongation observed in L248H cells. As mitochondrial elongation could result from an exacerbated fusion or inhibited fission, we suspected that these cells’ fission process is affected.

**Figure 3.**
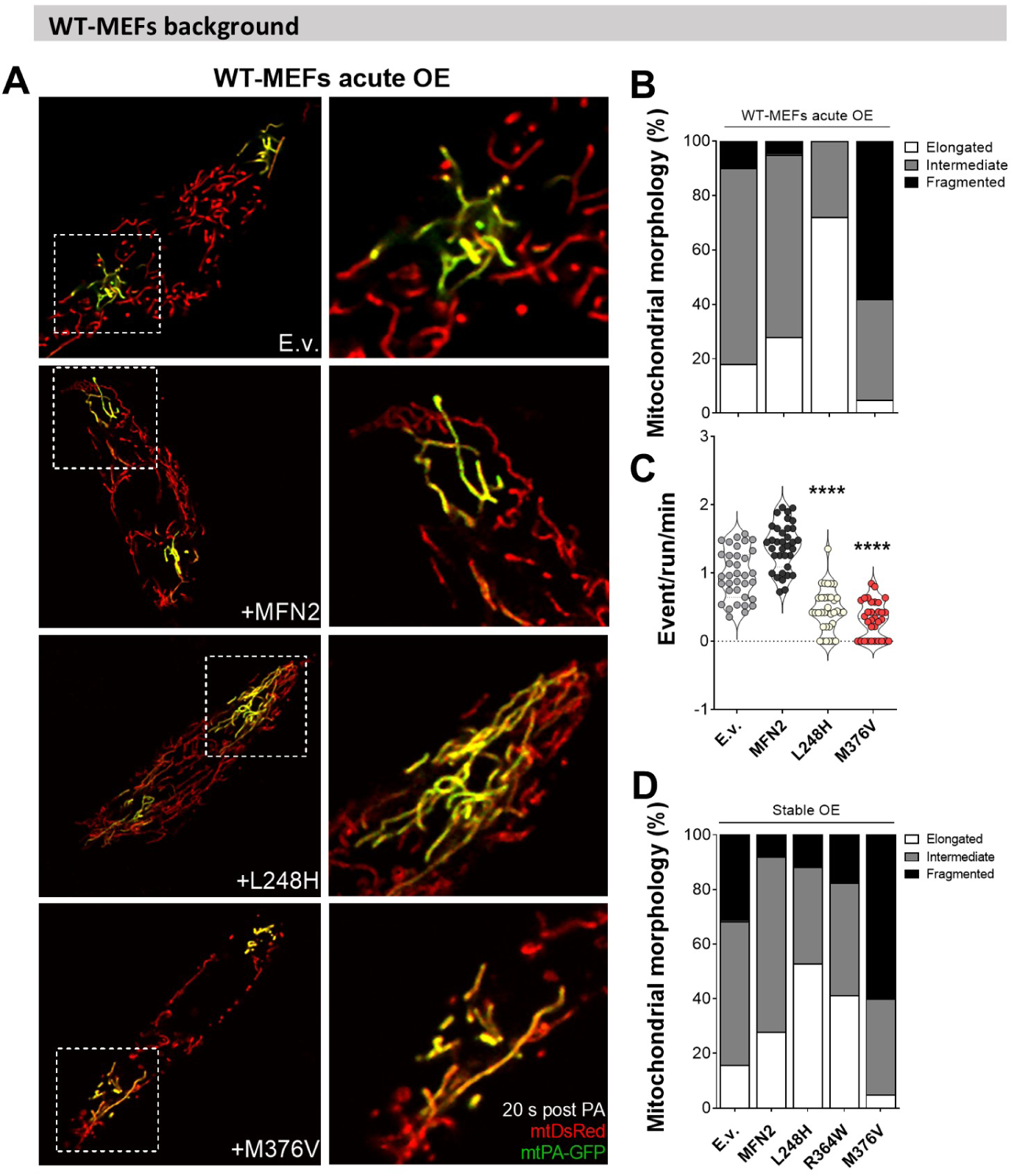
MEFs expressing CMT2A-causing mutants show low mitochondrial fusion frequency and differentially altered morphology. (**A**) WT-MEFs co-expressing mtDsRed, mtPA-GFP, and WT or mutant MFN2. Representative images 20 s post-photoactivation of 5×5 μm ROIs (left). Insets, displaying 1 ROI highlighting the extent of mtPA-GFP diffusion and mitochondrial morphology (right). (**B**) Mitochondrial morphology classification and quantification using mtDsRed images in ≥ 30 cells. (**C**) Fusion events frequency (MEF + E.v. = 35 cells, MEF + MFN2 = 35 cells, MEF + MFN2-L248H = 35 cells, MEF + MFN2-M376V = 32 cells). (**D**) Mitochondrial morphology classification and quantification using mtDsRed images in MEFs stable-expressing the E.v., WT MFN2, or mutants. Data were collected from ≥17 cell per condition (MEF + E.v.= 19 cells, MEF + MFN2 = 25 cells, MEF + MFN2-L248H = 17 cells, MEF + MFN2-R364W = 24 cells MEF + MFN2-M376V= 20 cells). Data are mean ± SEM of ≥ 3 independent experiments. ****P<0,0001 *vs* MFN2.

To evaluate the effect of chronic expression of CMT2A-mutants, we generated stable MEFs expressing WT or mutated MFN2-Flag (Fig. S2). Furthermore, we incorporated the mutant MFN2-R364W, which has been reported to cause hyperfusion of mitochondria ^32,33^. These cells recapitulate the abnormal morphology observed in the patient’s fibroblasts and WT MEFs upon acute overexpression model (Fig. 3D). The expression of the CMT2A-causing mutants did not alter the expression levels of endogenous Mfn2 (Fig. 4A-C) and correlates with mRNA levels measured by qPCR (Fig. 4D). The stable overexpression of WT MFN2, and MFN2 M376V significantly reduced the Mfn1 protein levels, however, cells expressing the mutation L248H or R364W, did not show any compensatory change on MFN1 levels (Fig. 4E-F), presumably due to the low expression levels of these particular mutations. Also, MFN2 overexpression induced an increase in long and short Opa1 protein levels (Fig. 4E). However, expression of the mutations L248H and R364W induced a change in the OPA1 processing with mostly short forms (Fig. 4E-G-H). No increase was observed in M376V-expressing cells (Fig. 4E-G). These results do not fully mimic the protein profiles observed in patients’ cells. However, our different WT background models maintained the morphological and dynamic features, suggesting that the differences in fusion protein levels are not the main drivers of the mitochondrial morphology phenotype.

**Figure 4.**
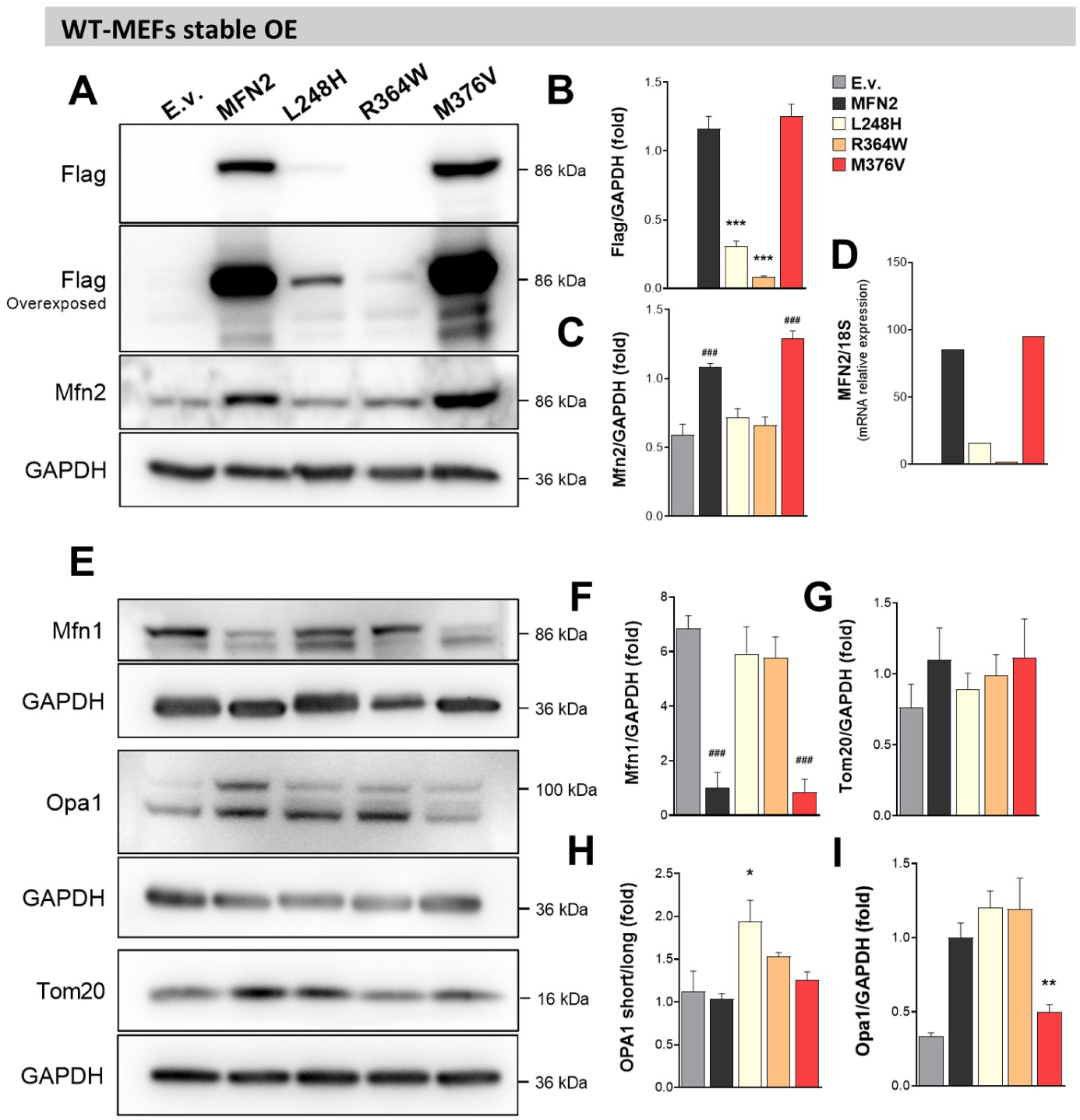
Effect of WT or mutant MFN2 stable expression on mitochondrial fusion proteins in WT MEFs. (**A**) and (**E**), Representative images from whole protein extracts Western blot of WT MEFs stable-expressing WT or mutants MFN2. (**B**), (**C**), (**F**), (**G**), (**H**), and (**I**) Western blot quantification from A and E panels. (**D**) MFN2 mRNA relative levels analyzed by RT-qPCR, N=1. Data are mean ± SEM of ≥ 3 independent experiments. ### P<0,001 vs E.v.; **P<0.01, ****P<0.0001 *vs* MFN2.

### Divergent morphological defects in L248H and M376 result from differential fission dynamics and Drp1 phosphorylation

#### Mitochondrial fission studies in CMT2A patient-derived fibroblasts

Our observation on mitochondrial morphology cannot be fully explained by the effect of MFN2 mutations on mitochondrial fusion; thus, we next tested if mitochondrial fission can also be involved. Previously, it has been proposed that mitochondrial fusion and fission machinery colocalize in the ER-mitochondria interface ^17^, suggesting that MFN2’s role in fusion could disturb fission. To further explore if fission dynamics were disturbed in patients’ cells carrying *MFN2* mutations, we evaluated mitochondrial fission as previously described ^11^. We quantified mitochondrial length, fission frequency, lag time, and number of constrictions per mitochondrial length (Fig. S3A, 5A). As expected, MFN2 L248H-carrying cells showed an increase in mitochondrial length when compared with fibroblast from control individuals, confirming our previous observation (Mean length: *Control 1*= 6.7 *μm*; *L248H* = 9.6 *μm*; *M376V* = 5.7 *μm*) (Fig. 5B). On the other hand, cells carrying MFN2 M376V mutations showed no difference as we consider only mitochondria longer than 2 µm for this analysis (*see methods*). Cell carrying the MFN2 L248H mutation showed low fission frequency and increased lag time (Mean Lag time *Control 1* = 55 s; *L248H*=93 s; *M376V*=33 s), together with a high number of unproductive mitochondrial constrictions sites (Fig. S3A-B). MFN2 M376V-carrying cells displayed an increased tendency in fission frequency; however, this was not significant when compared with control fibroblasts (Fig. 5C). All in all, these results are consistent with our observations for cells carrying the MFN2 L248H mutations with elongated mitochondrial, explained by impaired mitochondrial fusion and low mitochondrial fission rate, suggesting a role of this mutation in the control of both fusion and fission.

**Figure 5.**
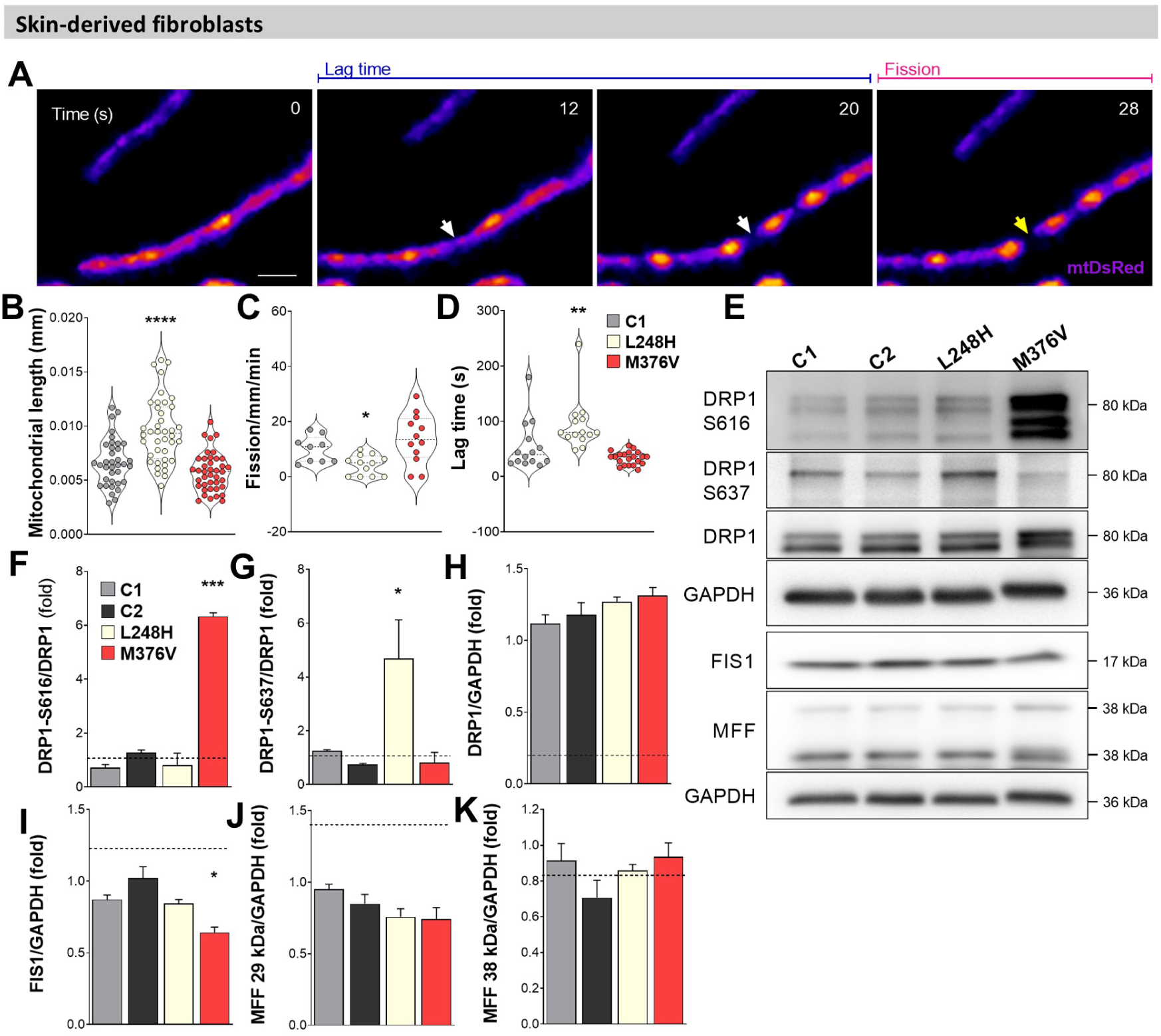
CMT2A patient-derived fibroblasts exhibit impaired mitochondrial fission dynamics and altered DRP1 phosphorylation status. (**A**) Control or CMT2A-derived fibroblasts expressing mtDsRed were imaged for 7 min. Fire LUT was applied to better visualize mitochondrial constrictions (white arrow). Image series were analyzed frame-by-frame, seeking fission events. Mitochondria separation into two units was defined as a fission event (Time 28 s, yellow arrow). The lag time corresponds to the time elapsed between the appearance of a constriction (Time 12 s) and the frame previous to the fission event at this site (Blue line). (**B**) Mitochondria length, (**C**) fission frequency and (**D**) Lag time were quantified in ≥40 mitochondria per condition from at least 10 different cells. (**E**) Representative Western blot from whole protein extracts of Controls (C1 and C2) or CMT2A-derived fibroblasts carrying MFN2 mutants. (**F**), (**G**), (**H**), (**I**), (**J**), and (**K**) densitometric quantification. The dashed line represents the C1 and C2 average. Data are mean ± SEM of ≥ 3 independent experiments. *P<0.05, **P< 0.01, ****P<0.0001 *vs* Control 1.

To test the involvement of the fission machinery proteins in the observed phenotypes, we studied specific DRP1 phosphorylation related to fission activation (p-Serine 616) and inhibition (Serine 637). Western blot analysis showed a distinct phosphorylation of DRP1 with increased levels of pSer616-DRP1 and pSer637-Drp1 in M376V and L248H, respectively (Fig. 5E-G). We also tested the protein levels of DRP1 anchors FIS1 and MFF and found diminished levels of FIS1 protein in M376V cells compared to controls (Fig. 5E-I). No changes were found in the MFF levels (Fig. 5E-J-K). Our data suggest that the differential variations observed in the mitochondrial morphology from patients’ cells carrying the MFN2 L248H and M376V mutations could be partially explained by the different Drp1 phosphorylation status between the two MFN2 mutations.

#### Fission dynamic studies in stable cells expressing CMT2A-MFN2 mutants

We further explored mitochondrial fission dynamics in stable MEFs expressing MFN2 mutations. The apparent separation of two mitochondria does not guarantee the fully executed fission event, as the splitting organelles could be tethered and not be fully separated into individual organelles. To overcome this, we co-transfected the cells with mtBFP and a fluorescent Drp1 to visualize its localization at the fission sites during the division of two mitochondria. Cells were imaged for 7 minutes and processed to improve the visualization of Drp1 puncta over the mitochondria during the fission events (Fig. S4A and S4B). Using this methodology, we verified that 100% of analysed fission events were marked with Drp1 in the fission site (Fig. 6A, S4A-B).

**Figure 6.**
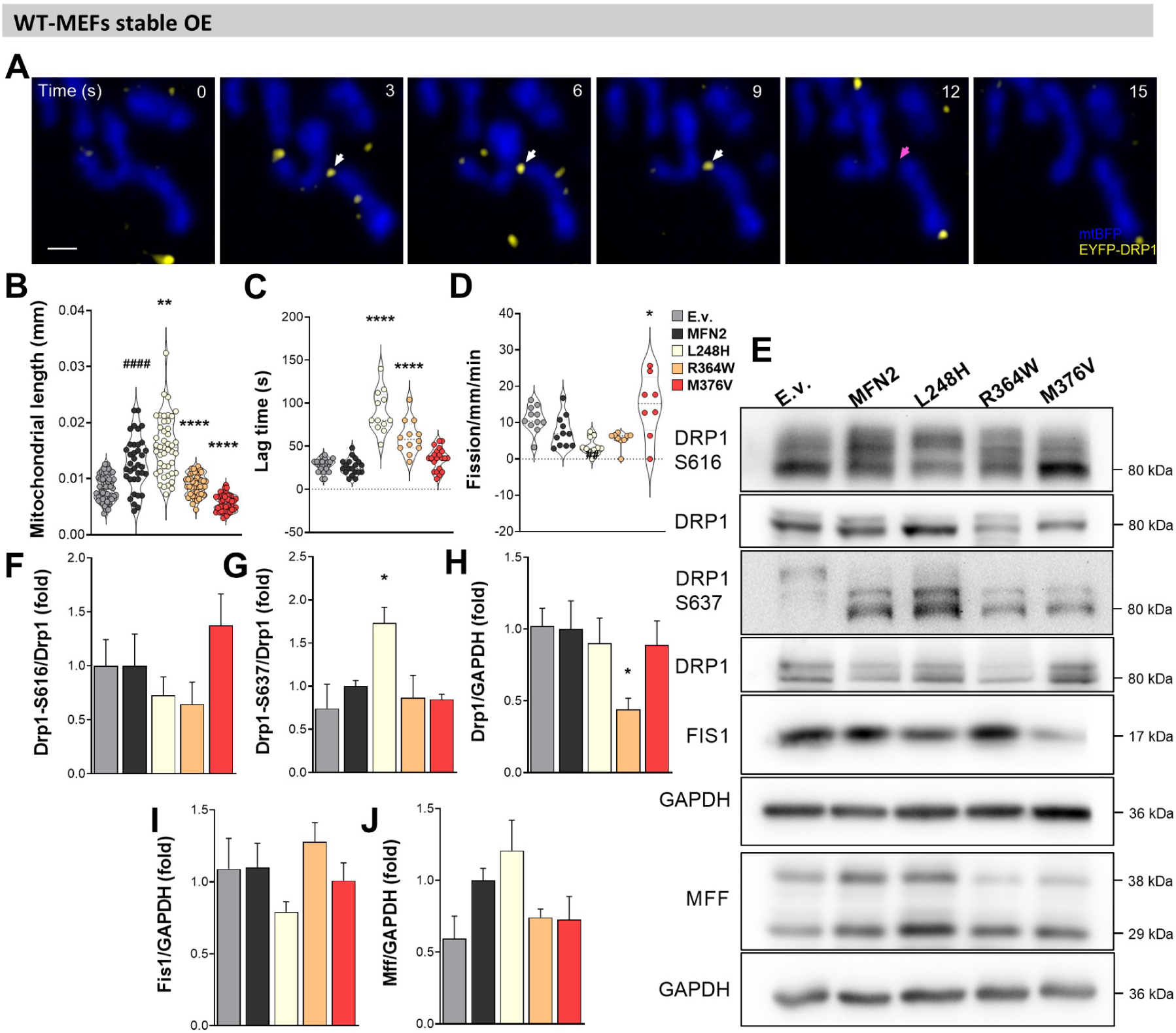
Hyperelonged-phenotype mutants MFN2-L248H and MFN2-R364W impair mitochondrial fission dynamics in stable MEFs. (**A**) MEFs stable-expressing MFN2 or mutants co-transfected with mt-BFP and EYFP-DRP1. EYFP-DRP1 localizes at mitochondria (white arrows) until the fission execution (pink arrow). (**B**) Mitochondrial length, (**C**) Lag time, and (**D**) fission frequency were quantified in ≥40 mitochondria per condition from at least 10 cells. (**E)** Representative images from whole protein extracts Western blot of WT MEFs stable-expressing WT or mutants MFN2. (**F**), (**G**), (**H**), (**I**) and (**J**) densitometric quantification from panels in E. ^##^*P*<0.01, ^####^P<0.0001 *vs* E.v. and \**P*<0.05, \*\**P*< 0.01, ****P<0.0001 *vs* MFN2.

MFN2 overexpression induces an increase in the mitochondrial length (Fig. 6B), probably as a result of increased fusion, as we distinguished no changes in lag-time or fission rate (Fig. 6C). In MEFs carrying the MFN2 L248H mutation, we detected increased length (Fig.6B), accompanied by longer fission time (Fig. 6C) and reduced fission frequency (Fig. 6D). Different fission alteration was observed upon R364W, as these cells show decreased mitochondrial length, compared to cells overexpressing MFN2, with increased lag-time and no significant decrease in fission frequency (Fig. 6B-D), suggesting a milder defect of fission dynamics in these cells. Finally, patient cells with M376V carry the smallest mitochondria among the evaluated MFN2 mutants, with no changes in time of fission execution but significantly elevated fission events frequency (Fig. 6B-D).

Our fission dynamics studies in stable cell lines show a defect in fission dynamics with decreased frequency in L248H and a mild defect in R364W accompanied by augmented lag time. These results also confirmed our observations in the patient’s cells and the potential role of MFN2 on mitochondrial fission since the MFN2 L248H mutation suppressed fusion and fission. Considering the difference in fusion activity between these mutants, as R364W has been syndicated as a “gain-of-function” mutant ^32^, we can suggest that the alterations in the fission balance might be in response to a different mechanism. Otherwise, results in M376V cells confirm increased fission frequency, highlighting a role in the fusion/fission crossroad.

Next, we evaluated the levels of proteins involved in fission. As in patients, we detected increased anti-fission pSer637-Drp1 in MFN2 L248H carrying cells; meanwhile, the pro-fission pSer616-Drp1 was elevated in cells expressing the MFN2 M376V (Fig. 5E-G), suggesting that the differential mitochondrial phenotype in these cells is a result of the specific phosphorylation status of Drp1, determined by the mutants presence. Differently, we observed decreased levels of total Drp1 in R364W stable MEFs but no changes in Drp1 phosphorylation (Fig. 5E-H) as previously reported^33^. Finally, we studied the levels of the Drp1 anchors Fis1 and Mff. Although no significant changes were seen in Fis1 by WT MFN2 or MFN2 L364W overexpression, we found a slight reduction in cells carrying the MFN2 mutations L248H and M376V (Fig. 6E, I). WT MFN2 and MFN2 L248H overexpression induced an increase in Mff protein levels. At the same time, no changes were observed in cells carrying the MFN2 R364W or M376V mutations (Fig. 5E, J), suggesting a compensatory increase upon the mitochondrial hyperelongation in these cells.

Altogether, these data indicate that the MFN2-disease-causing mutants display distinct fission dynamics associated with different molecular adaptations in fission proteins, especially in Drp1 pro and anti-fission phosphorylation status.

### CMT2A-causing mutants differentially alter mitochondrial metabolism

The primary function of mitochondria is to transform energy to synthesise ATP. Mitochondrial DNA encodes crucial subunits of OXPHOS, key for mitochondrial bioenergetic performance ^54^. Thus, we tested the mtDNA levels and integrity in stable cells expressing WT and mutant MFN2. Our results show that MFN2 overexpression induced downregulation of mtDNA levels, with no significant change between WT and mutant MFN2 (Fig. S5A). Next, long-range PCR showed no deletions in MFN2 stable cell lines (Fig. S5B).

We measured the OCR and extracellular acidification rate to further characterize the impact of MFN2 mutations on mitochondrial bioenergetics. Our results show that MFN2 expression did not change the OCR (Fig. 7A-B). Strikingly, cells stable expressing MFN2 L248H mutant showed increased basal OCR and maximal respiration (Fig. 7A-C) and ATP-linked and spare capacity (Fig. S6A, C). Cells expressing the MFN2 R364H mutations showed a less pronounced increase in basal and ATP-linked respiration, while cells expressing the MFN2 M376V mutation showed a decrease in basal and ATP-linked respiration (Fig. 7A-B). The oxygen not used in ATP synthesis was calculated based on proton leak, which was augmented in all MFN2 mutant conditions (Fig. S6B), whereas non-mitochondrial respiration increased in L248H and R364W expressing cells (Fig. S6D). ECAR is related to the production of lactate and H^+ 55^. From these data, it is possible to calculate the basal ECAR and the glycolytic capacity, represented as the difference between basal ECAR and ECAR measured after complex V inhibition through oligomycin addition. We found a marked increase in ECAR of cells expressing the M376V mutant and a discrete change in R364W (Fig. 7D and 7E). Overexpression of WT MFN2 increased glycolytic capacity regarding empty vector expression, while R364W and M376V show a slight and marked decrease, respectively (Fig. S6F), suggesting a lower metabolic flexibility. Finally, we found an increased resting mitochondrial membrane potential in the presence of all the evaluated mutants (Fig. 7F). These results suggest an increased OXPHOS activity in L248H and R364W that display a hyper-elongated mitochondrial morphology and a glycolytic-based metabolism in cells carrying the M376V mutation which showed fragmented mitochondria.

**Figure 7.**
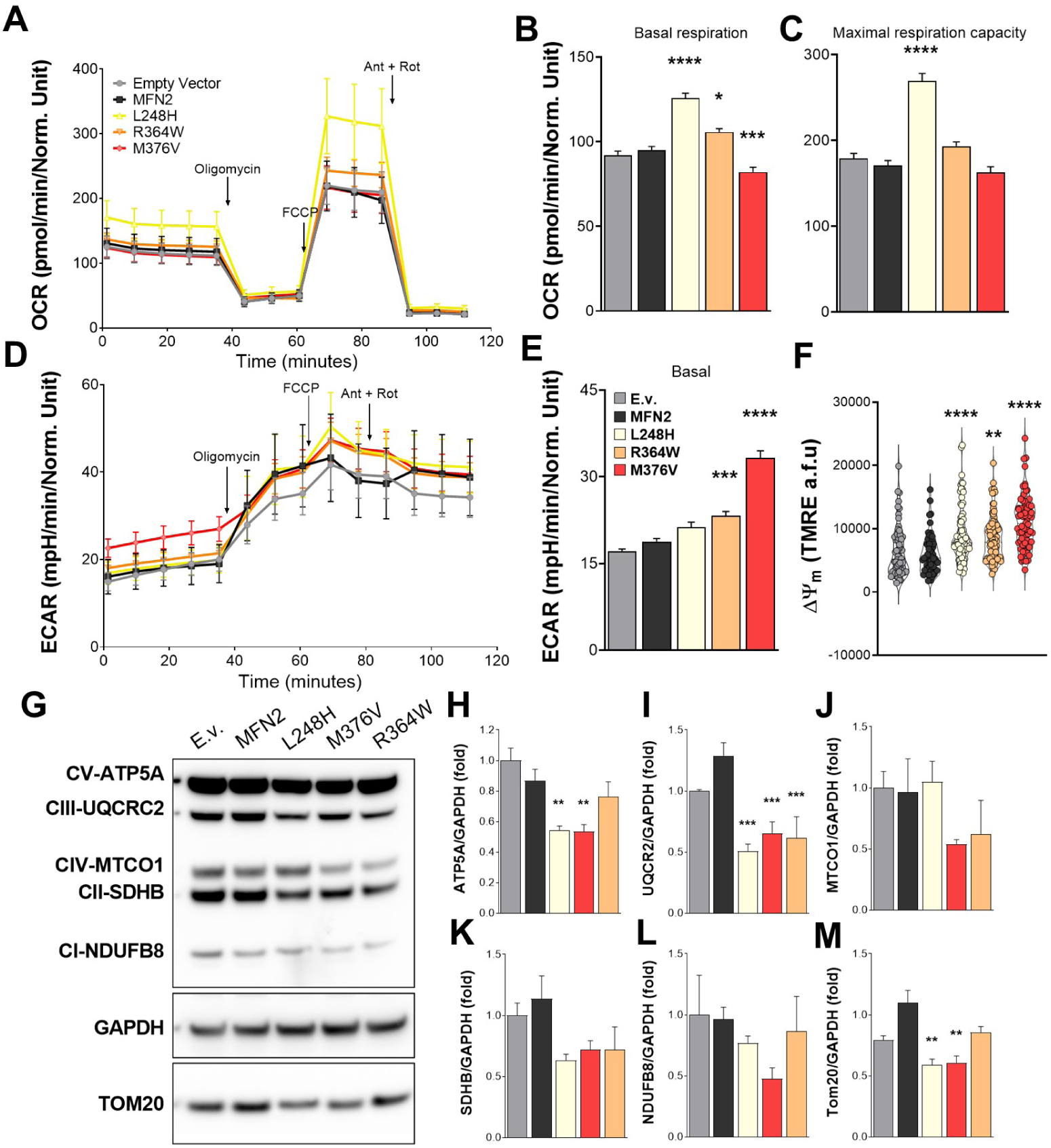
Stable MEFs expressing CMT2A-causing mutants display dissimilar oxygen consumption rates and OXPHOS alteration. (**A**) Representative oxygen consumption rates (OCR) and (**D**), extracellular acidification rate (ECAR) in stable MEFs expressing WT or mutants MFN2. OCR-based calculations are shown in (**B**), Basal respiration, and (**C**), Maximal respiration capacity. (**E)** Basal extracellular acidification rate (ECAR). (**F**) Resting mitochondrial membrane potential Δψm (*see methods*) (E.v., n = 3/58; MFN2, *n* = 3/58; L248H, *n* = 3/81; R364W, *n* = 3/93; M376V, *n* = 74). Data are mean ± SEM of ≥ 3 independent experiments. Data are mean ± SEM of ≥ 3 independent experiments. Seahorse experiments were performed using 12 technical replicates. (**G**) Representative Western blot of respiratory chain subunits from whole protein extracts from stable MEFs expressing WT or mutants MFN2. (**H**), (**I**), (**J**), (**K**), (**L**), and (**M**) densitometric quantification from panels in G. Data are mean ± SEM of ≥ 3 independent experiments. * *P*<0.05, ** *P*<0.01, *** *P*<0,001 **** *P*<0.0001 *vs* MFN2. *### P*<0,001 *vs* E.v.

To expand our understanding of changes in mitochondrial function upon MFN2 mutant expression, we studied OXPHOS subunit protein levels. We found reduced levels of complex V subunit ATP5A in stable cells expressing L248H and M376V and a non-significative reduction in R364W cells (Fig. 7G-H). Moreover, all the studied mutants showed downregulation of complex III subunit UQCRC2 and a substantial reduction tendency in complex II subunit SDHB (Fig. 7G, I, K). At the same time, complex IV subunit MTCO1 protein levels are not significantly reduced in M376V and R364W and maintained levels in MFN2-L248H (Fig. 7G and 7J). Finally, we found a slight reduction of the complex I subunit NDUFB8 protein levels in cells expressing MFN2 L248H and R364W and a more pronounced downregulation in MFN2 M376V cells, with no significant differences (Fig. 7G, L).

Overall, the results suggest that MFN2 mutants impair the bioenergetic metabolism, partly disrupting the levels of OXPHOS subunits independently of the mtDNA levels and stability. Differential levels of complex IV and I subunits might impact the bioenergetic differences between mutants.

## Discussion

Mutations in MFN2 are the primary cause of CMT2A ^21^. In this work, we show that MFN2 mutations perturb both mitochondrial fusion and fission dynamics by differential modification of fusion and fission machinery, with a particular impact on the DRP1 phosphorylation state, which tunes the mitochondrial morphology network to fragmentation or hyperelongation (Supplementary Table 4).

MFN2 mutations, the leading cause of CMT2A, result in various mitochondrial morphological changes.

Mild cristae ultrastructure defects were reported in the presence of CMT2A-MFN2 mutants ^48–50^. This evidence matches with the mild cristae alterations observed in our M376V fibroblasts. However, the severe cristae disruption observed in L248H cells is unprecedented, suggesting a differential cristae remodelling defect in the presence of this particular mutation. Patient-derived fibroblasts carrying different mutations in the GTPase domain (T105M, I213T, F240I, or V273G), or HR2 domain (L734V) show tubular morphology similar to control cells ^56^, while fragmentation was reported in motor neurons reprogrammed from patient fibroblasts bearing GTPase (T105M or R274W) or HR1 (H361Y or R364W) mutants, and in dorsal root ganglion neurons expressing mutants L76P, R94Q and P251A ^25,57^. In our model, cells carrying the MFN2 L248H mutation in the GTPase domain lead to hyperelongation, while cells carrying the mutation MFN2 M376V, located in the HR1 domain, lead to fragmentation. These data depict that mutant MFN2-mediated mitochondrial morphological changes strongly depend on the mutated amino acid and the cell type.

Mitochondrial hyperelongation is scarcely associated with CMT2A-causing mutation. For instance, hyperelongation was observed in transgenic flies expressing CMT2A-MFN2 mutants R364W and L76P, displaying hyper-fusion as a result of a gain of fusogenic activity and depletion of mitochondria at the neuromuscular junction ^32^, different to the hyperelongation with decreased fusion frequency that we observed in L248H cells. Thus, our results suggest that the mitochondrial hyperelongation observed in the cells carrying the MFN2 L248H mutation rises from the inhibition of both fusion and fission. In contrast, the fragmentation observed in the cells carrying the MFN2 M376V mutations results from decreased frequency of fusion and augmented fission events.

In our MFN2-KO model, the expression of WT MFN2 completely rescued mitochondrial morphology into an intermediate phenotype. Similarly, the expression of MFN2 L248H showed mainly an intermediate morphology, with a small proportion of hyper-elongation in L248H-carrying cells, while MFN2 M376V induces a partial rescue (Fig. S1B). Despite this morphology rescue, fusion frequency was decreased in the presence of both, consistent with the observations in patient cells. These findings point out that mitochondrial morphology is not always directly correlated with the integrity of mitochondrial fusion dynamics, and more specialized studies, beyond morphometric analysis, are necessary to conclude about alterations in mitochondrial shape and the processes influencing it. Different authors proposed that MFN2 mutants can exert a “dominant-negative” effect, altering the function of the WT allele ^52,58,59^. This idea is consistent with our observations in our results in WT-MEFs cells expressing CMT2A-MFN2 mutants, which completely recapitulates the dynamic and morphological changes observed in patient fibroblasts in the presence of WT-MFN2 alleles. The molecular mechanism supporting MFN2-mediated fusion is not fully understood. Structural studies showed that homodimers (MFN2-MFN2) and heterodimers (MFN2-MFN1) formation depends on the interaction between the GTPase domains ^3^. Also, HR1 and HR2 intra-molecular and inter-molecular interactions are proposed as mediators of mitochondrial docking and fusion ^60^. Thus, intra and inter-protein interactions are relevant in determining MFN2 function and activity. The crystal structure of a portion of MFN2 lacking amino acids 1–21 and 401–705 (MFN2-IM) reveals that most CMT2A-causing mutants localize on the protein surface. Dimerization studies show that mutations MFN2-L248V and MFN2-R364W disrupt WT MFN2-mutant MFN2-IM interaction, while no changes were found regarding MFN2-M376L mutant. Also, MFN2-L248V and MFN2-R364W lose the capacity to form homodimers (MFN2-IM – MFN2-IM) but not MFN2-M376L. Furthermore, these three mutants can form hetero-dimers with MFN1 ^3^. Thus, mutations in these residues may impair the interacting equilibrium of MFN2 in the mitochondrial fusion spot. Yet, our studies suggest that these mutants also disturb the mitochondrial fission machinery.

Mitochondrial fusion and fission co-exist as balanced processes occurring in a specialized niche, ensuring the execution of one or the other ^16,17,19^. Whether MFN2 participates in mitochondrial fission regulation and if fission defects may play a role in CMT2A disease is unknown. Our data reveal that hyperelongation in patients’ cells or MEFs carrying L248H is a result of dampened fission dynamics along with augmented levels of DRP1-pS637, known for promoting fission inhibition with increased mitochondrial constrictions ^61^, in agreement with our observations. Heterozygous missense mutations of DRP1 produce hyperelongation and cause diseases with neuromuscular phenotypes in patients ^62^, illustrating the deleterious effects of this phenomenon in humans. Interestingly, DRP1 and MFN2 share different regulators, such as kinases and ubiquitin ligases, such as PKA^63,64^, PINK1 ^65,^ and MITOL ^66^, but how this cross-regulation occurs is still unknown. To our knowledge, this is the first report showing an effect on DRP1 phosphorylation status due to MFN2 mutants.

MFN2-R364W was described as a gain of function mutation increasing mitochondrial fusion ^32,33^. Our results show a mild impairment in fission dynamics, a non-significant increase in lag time, and decreased levels of total Drp1. Recent data show that expression of R364W in HeLa cells induces a reduction in Drp1 levels caused by augmented ubiquitylation via the E3-ubiquitin ligase MITOL and proteasomal degradation ^33^. Thus, L248H and R364W are likely to cause mitochondrial hyper-elongation by different mechanisms. Finally, we found that M376V patient-derived and stable cell cells exhibit increased mitochondrial fission frequency. Thus, the mitochondrial fragmentation observed in M376V-expressing cells may result from increased underlying fission and inhibited fusion in these cells. Also, these results suggest possible facilitation of fission activity, as supported by increased levels of pro-fission phosphor-DRP1-Ser161 protein in our study in patient fibroblasts and stable cell lines. DRP1 phosphorylation at Ser616 is associated with an increase in mitochondrial fragmentation as a result of the action of different kinases such as PINK1 ^65^. We hypothesize that MFN2 might interact with upstream kinases that regulate the activity of DRP1, and mutations may alter this crosstalk ^67^. Further experiments should focus on elucidating how MFN2 mutants impact the activity of DRP1.

The abundance and integrity of mtDNA determine the expression of thirteen OXPHOS complexes subunits ^68^. In the MFN2-overexpressing model, we found decreased mtDNA levels without integrity compromise. Low mtDNA levels were reported in different models lacking MFN2 ^69–71^; however, mutations promote higher mtDNA copy numbers in the blood of patients ^72^, suggesting a complex interplay between disturbed mitochondrial dynamic and mtDNA stability. No data regarding the effect of MFN2 overexpression on mtDNA levels are available. The replication of mtDNA is coupled to ER-contact sites and mitochondrial division, a mechanism that ensures the inheritance of mtDNA into the nascent mitochondria^13^. Also, the localization of mtDNA through mitochondria is marked by ER, and it is proposed to be mediated by MIRO/KIF5B/Mic60 ^73^. Thus, taking into account that i) the MFN2 overexpression might cause miss-localization of the protein on the OMM, ii) MFN2 is one of the ER-mitochondria tethers that support the contacts ^74^, and iii) it participates in the mitochondrial transport interacting with Miro1 ^24^, we speculate that overexpression of WT MFN2 or the mutants, might impair mtDNA replication and maintenance, by mechanisms linked to mtDNA mislocalization.

New insights show how MFN2 modulation impacts mitochondrial function, using small molecules targeting the interaction of the HR1 and HR2 domain, which is crucial to determining pro or anti-fusion conformation^75^. Pro-fusion molecule drives elongation of mitochondria with increased respiration and membrane potential, as we also observed in our hyperelongated mutant, MFN2 L248H, although our phenotype was determined by fission inhibition rather than increased fusion. Thus, in our model, hyperelongation seems detrimental, likely contributing to the pathomechanism of the L248H mutation in CMT2A. Moreover, we found that cells carrying MFN2 M376V are less oxidative in basal conditions and more glycolytic, as our increased ECAR results suggest. These results are in agreement with the observed MFN2 knockdown cell that shows reduced mitochondrial respiration ^76^, and with results in permeabilized muscle fibers and fibroblasts from patients carrying the M376V displaying a reduction in respiration rate with round mitochondria and cristae affectation in muscle ^77^. Strikingly, our data showed increased membrane potential. However, ATP-linked respiration was dampened, and OXPHOS subunit levels were altered, suggesting a possible increase in the reverse activity of complex V, consistent with previous reports under glycolytic conditions ^78^.

In conclusion, we show that CMT2A-MFN2 mutations L248H and M376V inhibit mitochondrial fusion and differentially disrupt the mitochondrial fission dynamics and oxidative metabolism, showing for the first time that MFN2 mutations may modulate mitochondrial fission in a pathogenic context, stressing that future pharmacological approaches focused on mitochondrial dynamics modulation in CMT2A, need to consider mutant-specific effects on mitochondrial fusion and fission processes.

## Data availability

The authors confirm that the data underpinning the conclusions drawn in this study can be found both in the main article and its Supplementary material.

## Acknowledgements and Funding

We thank Heidi McBride for the plasmid backbone encoding MFN2-ECFP plasmids and György Hajnóczky for helpful discussion. We thank Dr Enrique Brandan for his help with reagents and equipment. Finally, we thank the Chilean Government supported this work through Agencia Nacional de Investigación y Desarrollo (ANID) PhD fellowship 21191304 to DL, 21181402 to BC-S, Vicerrectoria de Investigación Pontificia Universidad Católica de Chile (VRI-UC) PhD fellowship to DL and BC-S. RH is supported by the Wellcome Discovery Award (226653/Z/22/Z), the Medical Research Council (UK) (MR/V009346/1), the Addenbrookes Charitable Trust (G100142), the Evelyn Trust, the Stoneygate Trust, the Lily Foundation, Ataxia UK, Action for AT, the Hereditary Neuropathy Foundation, the Muscular Dystrophy UK and the LifeArc Centre to Treat Mitochondrial Diseases (LAC-TreatMito). She is also supported by an MRC strategic award to establish an International Centre for Genomic Medicine in Neuromuscular Diseases (ICGNMD) MR/S005021/1. This research was supported by the NIHR Cambridge Biomedical Research Centre (BRC-1215-20014). The views expressed are those of the authors and not necessarily those of the NIHR or the Department of Health and Social Care. The study was supported by FONDECYT grants 1150677, 1191770 and 1231557 to VE.

---

**Figure S1. CMT2A-causing mutants cannot rescue the mitochondrial fusion frequency and provoke mild defects in morphology upon acute overexpression in Mfn2-/- MEFs.** (**A**) Mfn2-/- MEFs acutely co-expressing mtDsRed, mtPA-GFP, and WT or mutant MFN2. Representative images 20 s post-photoactivation of 5×5 μm ROIs (left). Insets displaying 1 or 2 ROIs highlighting the extent of mtPA-GFP diffusion and mitochondrial morphology (right). (**B**) Mitochondrial morphology classification and quantification using mtDsRed images in ≥ 30 cells. (**C**) Fusion events frequency (Mfn2-KO + E.v. = 19 cells, Mfn2-KO + MFN2 = 17 cells, Mfn2-KO + MFN2-L248H = 17 cells, Mfn2-KO + MFN2-M376V = 22 cells). (**D**) Representative Western blot from whole protein extracts of MEFs Mfn2-KO cells acutely expressing MFN2 WT or mutants. Data are mean ± SEM of ≥ 3 independent experiments. **P< 0.01 *vs* MFN2.

**Figure S2. MEFs stable-expressing CMT2A-causing mutants.** (**A**) Representative Western blot from whole protein extracts of MEFs clones stably-expressing WT or mutant MFN2. Clones highlighted in red were selected for the following experiments. (**B**) Selected clones were incubated for 20 min with TMRE (10 nM). Representative images and insets depicting mitochondrial morphology (dashed squares).

**Figure S3. Cells expressing MFN2-L248H show a bead-on-string mitochondrial phenotype.** (**A**) Representative images of Control or CMT2A-derived fibroblasts expressing mtDsRed showing mitochondrial matrix constriction (red arrowheads) and the fluorescence histogram through the mitochondrion. The red arrow point pronounces constriction sites. (**B**) Number of mitochondrial matrix constrictions per mitochondrial length (µm) in human fibroblasts and (**C**) stable MEFs.

**Figure S4. EYFP-DRP1 image pre-processing**. (**A**) MEFs stable-expressing MFN2 or mutants, co-transfected with mt-BFP and EYFP-DRP1, were imaged every 3 s for 7 min upon 405 and 514 nm laser excitation. The threshold was defined according to the integrated density average of 5 high-intensity EYFP-DRP1 puncta (bottom, white circles). (**B**) Mitochondrial fission example where EYFP-DRP1 high-intensity puncta (right, green arrowheads) concur with mitochondrial constriction (left, histograms).

**Figure S5. MFN2 overexpression decreases mtDNA levels in stable MEFs**. Total DNA extracted from stable MEFs expressing WT MFN2 or mutants. (**A**) The mtDNA/nDNA ratio was assessed using qPCR targeting the mitochondrial ND5 gene and the nuclear β-actin gene. (**B**) Agarose gel of three different experiments (n1, n2, and n3) showing long-PCR products of mtDNA amplification using the following primers: **mtDNA 7.7 kb F** ACCTGAATTGGGGGCCAACC; **mtDNA 7.7 kb R** TGGCGAAGTGGGCTTTTGCT and **mtDNA 8.6 kb F** AGCAAAAGCCCACTTCGCCA; **mtDNA 8.6 kb R** GGTTGGCCCCCAATTCAGGT. Data are mean ± SEM of ≥ 3 independent experiments. \*\**P*<0.01, \*\*\**P*<0,001 *vs* E.v.

**Figure S6. Seahorse OCR-based calculations.** OCR-based calculations representing (**B**) respiration linked to ATP production, (**B**) Spare capacity, (**C**) Proton leak, and (**D**) Non-mitochondrial respiration were plotted. (**E**) Coupling efficiency = (OCR ATP production)/(OCR Basal Respiration) x100; each consisting of 12 technical replicates. (**F**) Glycolytic capacity, calculated as the difference between ECAR following the injection of 1 μm oligomycin, and the basal ECAR. Data are mean ± SEM of ≥ 3 independent experiments. * P<0.05, **P<0.01, ****P<0.0001 vs +MFN2.

